# A Gender-Dependent Molecular Switch of Inflammation *via* MyD88/Estrogen Receptor-alpha Interaction

**DOI:** 10.1101/255778

**Authors:** Rana El Sabeh, Mélanie Bonnet, Katy Le Corf, Kevin Lang, Alain Kfoury, Bassam Badran, Nader Hussein, François Virard, Isabelle Treilleux, Muriel Le Romancer, Serge Lebecque, Serge Manié, Isabelle Coste, Toufic Renno

## Abstract

Most Toll-like receptors and IL-1/IL-18 receptors activate a signaling cascade via the adaptor molecule MyD88, resulting in NF-κB activation and inflammatory cytokine and chemokine production. Females are less susceptible than males to inflammatory conditions, presumably due to protection by estrogen. Here we show that MyD88 interacts with a methylated, cytoplasmic form of estrogen receptor-alpha (methER-α). This interaction is required for NF-κB transcriptional activity and pro-inflammatory cytokine production, and is dissociated by estrogen. Importantly, we show a strong gender segregation in gametogenic reproductive organs, with MyD88/methER-α interactions found in testicular tissues and in ovarian tissues from menopausal women, but not in ovaries from women age 49 and less -suggesting a role for estrogen in disrupting this complex *in situ*. Collectively, our results indicate that the formation of MyD88/methER-α complexes during inflammatory signaling and their disruption by estrogen may represent a mechanism that contributes to gender bias in inflammatory responses.

## Introduction

Inflammation is a biologically complex mechanism that aims to restore tissue homeostasis following various insults. In response to different triggers such as infection and trauma, the acute inflammatory response is induced with the production of various mediators that will act on the effector cells and tissues to eliminate the insult and to restore their homeostatic set points. This is followed by a resolution phase where tissue repair takes place. The persistence of the insult or an excessive immune response drive the inflammation from an acute to a chronic state that underlies several diseases.

It has long been observed that females are relatively protected from certain diseases with an underlying inflammatory component, such as atherosclerosis (1), ischemic stroke (2), neonatal hypoxic-ischemic encephalopathy (3), apidocyte atrophy (4), blood-brain barrier disruption (5), LPS-induced endotoxic shock (6), colitis (7), and cancers that develop on an inflammatory background, such as hepatocellular carcinoma (HCC) (8). The relative protection from inflammation is thought to be mainly due to the high levels of estrogen in pre-menopaused women (9-12). Menopaused or ovariectomized females have increased inflammation, whereas estrogen-treated males had milder inflammatory responses (13). Estrogen is produced both in males and females, but at much higher levels in the latter until menopause. Estrogen has been shown to be an inhibitor of inflammation, mainly *via* decreasing NF-kB signaling, leading to decreased cytokine and chemokine production (14, 15). In a model of liver inflammation leading to hepatocellular carcinoma, it was found that MyD88-dependent activation of interleukin-6 is negatively regulated by estrogen via the estrogen receptor alpha (ER-α) (16).

MyD88 was originally discovered as an adaptor protein in TLR and IL-1R signaling that induces the activation of the NF-kB transcription factor (17). MyD88 consists of an N-terminal death domain (DD) that enables MyD88 interaction with members of the IRAK kinase family and a C-terminal Toll-interleukin 1 receptor (TIR) domain that allows its interaction with TLRs (17).

Here we describe a novel mechanism of inflammatory signaling regulation via the interaction between MyD88 and a cytoplasmic form of ER-α, with implications for gender disparity in inflammatory responses.

## Results and Discussion

MyD88, a key adaptor protein in TLR/IL-1R signaling, harbors an evolutionarily conserved LXXLL motif in its death domain (Figure S1). This motif is necessary and sufficient for interaction with nuclear receptors (18). Since MyD88 and the nuclear receptor ER-α are functionally linked in inflammatory signaling (16), we asked whether these two proteins physically interact. As seen in Figure 1A, wild-type MyD88, but not forms of the protein truncated for the DD (ΔDD) or mutated in the LXXLL motif, imunoprecipitates with ER-α.

**Figure 1:**
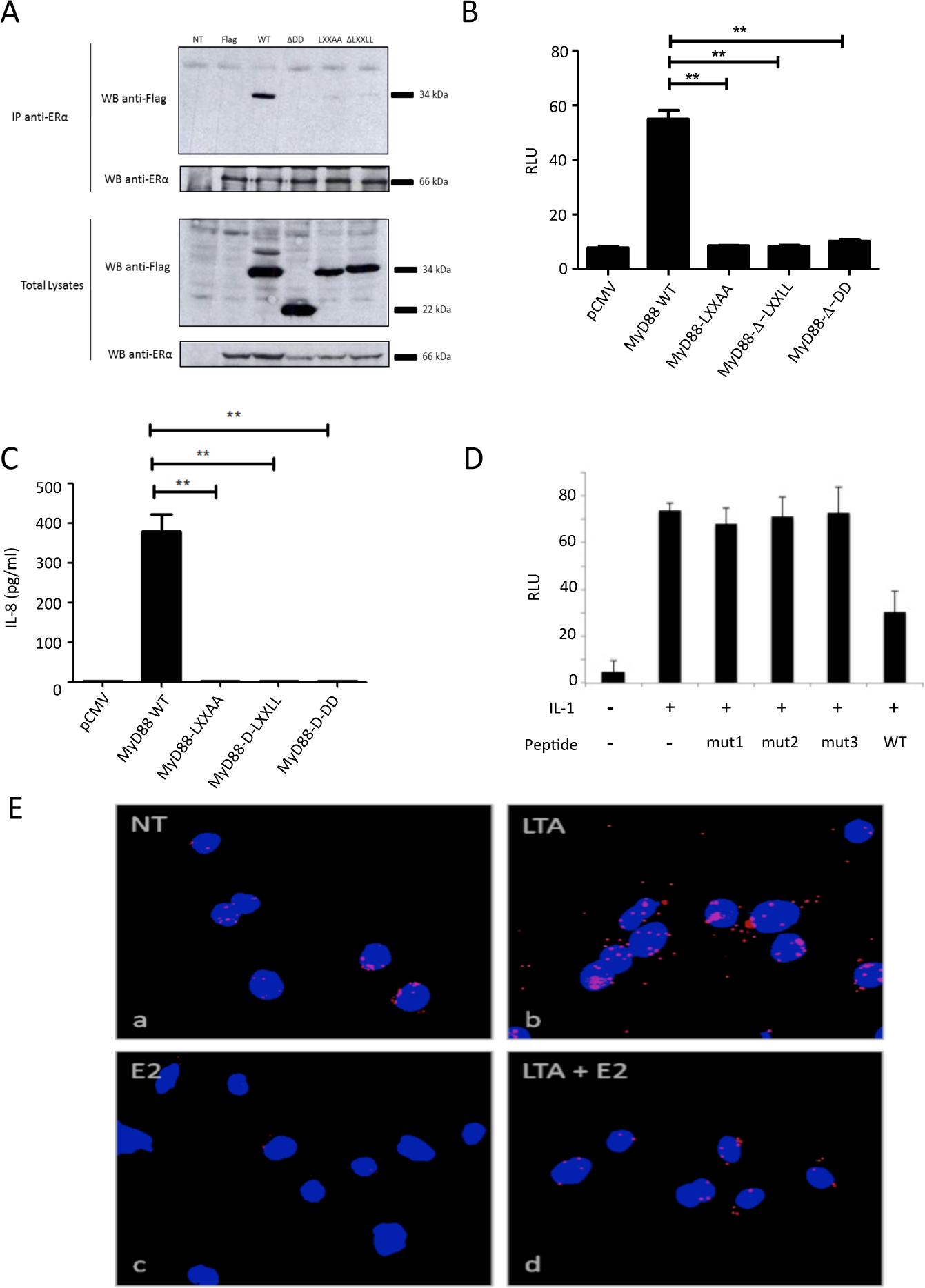
MyD88 forms a functional complex with ER-α. (**A**) HEK293T cells were transfected with a plasmid encoding ER-α and/or Flag-MyD88 (WT), Flag-MyD88-LXXAA (LXXAA), Flag-MyD88-delta LXXLL (ΔLXXLL), and Flag-MyD88-delta death domain (ΔDD). Lysates were immunoprecipitated with anti-ER-α antibody and immunoblotted with anti-Flag antibody. (**B**) Luciferase activity from NF-κB luciferase in HEK293T cells overexpressing MyD88 WT, LXXAA, LXXLL, or ΔDD. All luciferase values were normalized based on Renilla luminescence. (**), *P*<0.01.(**C**) IL-8 production by HEK293T cells was measured by ELISA in culture supernatants after transfection with MyD88 WT, MyD88-LXXAA, MyD88 ΔLXXLL, and MyD88-ΔDD plasmids. (**), *P*<0.01. (**D**) Luciferase activity from NF-κB luciferase in MCF7 cells, which were pretreated with cell-permeable peptides containing the LXXLL motif (WT) or a mutant form (mut), then stimulated with IL-1. (**E**). THP-1 monocytic cells were stimulated with LTA and/or estrogen (E2), then stained for MyD88 and ER-α according to the Duolink protocol. NT: non-treated cells. Magnification: 40x

To address whether this interaction plays a role in NF-κB activation, a NF-κB-luciferase reporter assay was performed, in which HEK293T cells were transfected either with wild-type, or with ΔDD or LXXLL mutated forms of MyD88 (Figure S2). Again, only wild-type MyD88 can activate NF-κB, whereas MyD88 mutant forms that cannot interact with ER-α are unable to do so (Figure 1B). As shown in Figure 1C, whereas wild-type MyD88 induces high levels of IL-8, the MyD88 mutant forms unable to interact with ER-α produced virtually no IL-8. To unequivocally ascertain the importance of MyD88/ER-α interaction itself to inflammatory signaling, we used a cell-permeable peptide (Figure S3) to inhibit the endogenous MyD88/ER-α interaction in IL-1-treated cells. This peptide contains the sequence of the MyD88 death domain containing the LXXLL motif responsible for the ER-α binding. As shown in Figure 1D, the specific peptide (WT), but not peptides that were mutated in the LXXLL motif (mut1-3), reduced NF-kB transcriptional activity in response to IL-1. Therefore, MyD88 interacts with ER-α via its LXXLL motif, and this interaction is required for NF-κB activation and cytokine production.

We then investigated the regulation of endogenous MyD88/ER-α interaction. To this end, we used the highly sensitive *in situ* proximity ligation assay (19) on the acute monocytic leukemia THP-1 cell line that is known to be responsive to both estrogen and TLR ligands (20, 21). THP-1 cells were treated with LTA (lipoteichoic acid, TLR2 agonist) and/or with estrogen, before performing PLA. As seen in Figure 1E and Figure S4, untreated THP-1 cells show some interaction between MyD88 and ER-α. However, upon treatment with LTA, there is an increase in the interaction between MyD88 and ER-α, which is strongly reduced in presence of estrogen. Therefore, MyD88 and ER-α interact at the endogenous level. This interaction is enhanced by TLR stimulation and is dissociated in the presence of estrogen.

ER-α signaling can be divided into two major pathways, the genomic and the non-genomic pathways. In the genomic pathway, ligand-bound ER-α translocates to the nucleus where it activates target gene transcription. The non-genomic or “rapid signaling” pathway is essentially cytoplasmic and is mainly mediated by methylated ER-α. Upon rapid estrogen stimulation (less than 5 minutes), the methyltransferase PRMT1 methylates ER-α on arginine 260 present in its DNA binding domain and which also lies within one of the nuclear localization signals of ER-α, thus maintaining methylated ER-α (methER-α) exclusively in the cytoplasm (22). Since MyD88 is also a cytosolic protein, we investigated whether it particularly interacts with methER-α. We showed in co-immunoprecipitation experiments that upon cell stimulation with IL-1, ER-α is methylated and it interacts with MyD88. Treatment of cells with IL-1 in presence of estrogen, or with siRNA specific for PRMT1, inhibits ER-α methylation, and therefore prevents ER-α interaction with MyD88 (Figure 2A, Figure S5).

**Figure 2:**
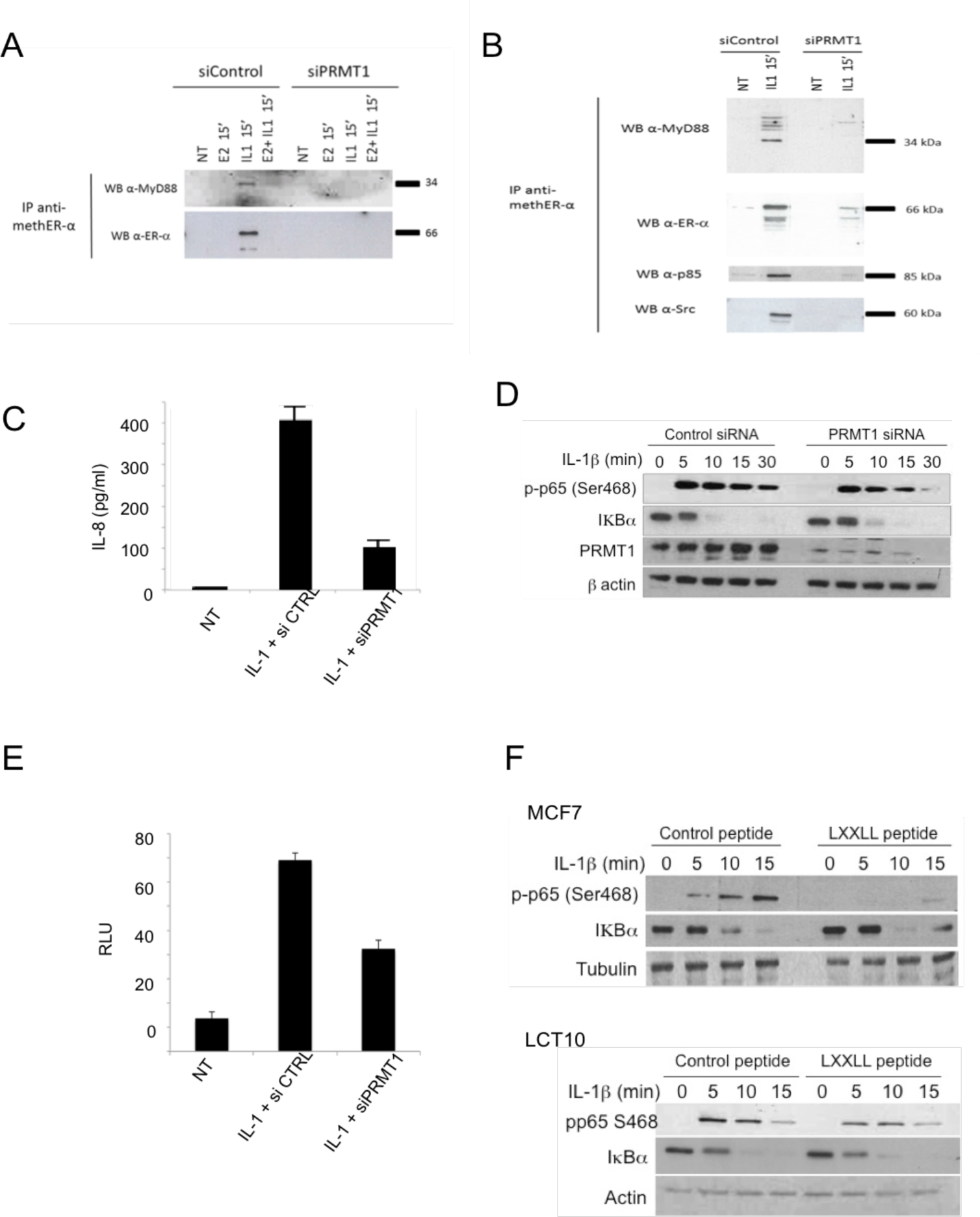
IL-1 stimulation leads to the formation of MyD88/methER-α complexes, which are required for NF-κB transcriptional activity and IL8 production. (**A**) MCF 7 cells were transfected with control siRNA (siControl) or PRMT1-specific siRNA (siPRMT1) and then treated with E2, IL-1, or both. Methylated ER-α was then immunoprecipitated and blotted with anti-MyD88 and anti-ER-α antibodies. (**B**) MCF 7 cells were transfected with siControl or siPRMT1 and were treated with IL-1. Methylated ER-α was then immunoprecipitated and blotted for MyD88, ER-α, p85, and Src. (**C-E**) MCF 7 cells were transfected with siControl or siPRMT1, and treated with IL-1. After 24h, IL-8 production was measured in culture supernatants by ELISA (**C**); the amount of p-p65(Ser468), IKBα, PRMT1 and actin was determined by western blot **(D**); and luciferase activity from an NF-κB-luciferase reporter was measured (E). Values were normalized based on Renilla luminescence. (**F**) MCF7 and LC10 cells were pretreated with cell-permeable WT or mutant (mut) peptides for 1h and then stimulated with IL-1 for the indicated times. The amount of p-p65(Ser468), IKBα, PRMT1, and actin was determined by western blot.

It has been shown that upon its methylation, ER-α interacts with p85, the regulatory subunit of PI3K, and with Src (22). To investigate whether MyD88, methylated ER-α, and these proteins are in the same complex, we performed an endogenous immunoprecipitation of methylated ER-α from MCF 7 cells upon IL-1 stimulation in the presence or absence of PRMT1. As shown in Figure 2B and Figure S6, we detected the presence of p85 and Src in the MyD88/methER-α complex under the same conditions. This is consistent with the findings that showed that MyD88 and p85 can be found in the same complex (23). These interactions were abrogated upon PRMT1 knockdown. Taken together, these data suggest that inflammatory signals trigger methylation of ER-α and its interaction with MyD88 in a complex containing PI3K p85 and Src.

To investigate whether the disruption of the MyD88/metER-α complex by PRMT1 knockdown is associated with biological consequences, MCF7 cells were stimulated with IL-1 after non-silencing or PRMT1-specific siRNA transfection. We found a profound decrease in IL-8 production when PRMT1 is silenced (Figure 2C, Figure S7). Since MyD88 is part of the IL-1 signaling machinery, we asked whether its interaction with methER-α depends on its function as an adaptor of IL-1R. We used the NCI-H292 lung cancer cell line, which responds to both IL-1R and TLR3 ligands by producing IL-8 *via* the MyD88 and TRIF adaptors, respectively. We found that knocking down PRMT1 inhibits IL-1- and Poly(I:C)-induced IL-8 secretion to a similar extent (Figure S8), suggesting that MyD88/methER-α is required for both MyD88-dependent and -independent inflammatory signaling.

We then sought to decipher the mechanisms whereby PRMT1-methylated ER-α and its association with MyD88 are implicated in the inflammatory response induced by IL-1. Since PRMT1 has been described to methylate mainly late downstream targets such as histones, transcription factors, and cell-cycle and repair proteins (24) and given that ER-α methylation by PRMT1 occurs early in the response to IL-1 (15 min, Figure S9), we reasoned that analyzing early consequences of PRMT1 silencing should reduce the contamination of the results with off-target, indirect effects.

With this in mind, we looked at NF-B activation in presence or absence of PRMT1. We found that early cytoplasmic events after IL-1 stimulation such as IκB degradation were not affected by knocking down PRMT1 (Figure 2D). At this time, most of the MyD88-methER-α complexes have not yet been formed (Figure S9). We therefore asked whether p65 translocation into the nucleus also occurs in absence of PRMT1. We showed that the p65 sub-unit of NF-B complex translocates into the nucleus between 15 and 30 minutes after IL-1 stimulation independently of PRMT1 expression levels (Figure S10). We then asked whether the p65 that was translocated into the nucleus in absence of PRMT1 is capable of activating the transcription of NF-B target genes. To address this question, a NF-κB-luciferase reporter assay was performed under PRMT1 inhibition. Results show that transcriptional NF-κB activity is reduced as early as 2 hours after stimulation in absence of PRMT1 (Figure 2E, Figure S7). A post-translational modification of p65, the serine 468 phosphorylation, had been described to occur in the nucleus 15 min after activation and to be essential for enabling NF-kB transcriptional activation of several target genes (25). We therefore assayed p65 S468 phosphorylation in response to IL-1 and found that this nuclear post-translational modification is reduced post-treatment with IL-1 when PRMT1 is knocked down (Figure 2D).

To ascertain whether lack of MyD88/ER-α interaction resulting from knocking down PRMT1 is responsible for the defect in p65 activation, we used the cell-permeable LXXLL peptide described above to inhibit the endogenous MyD88/ER-α interaction in IL-1-treated cells. Similar to what was observed upon PRMT1 knockdown, the LXXLL-containing peptide did not affect IkB degradation (Figure 2F). However, also as with siPRMT1, the specific peptide inhibited p65 phosphorylation on S468 in response to IL-1 in both MCF7 cells and the mouse Leydig cancer cell line LCT10 (Figure 2F).

To examine whether MyD88/methER-α interaction also occurs in human tissue, we performed PLA on tissue microarrays with a wide panel of tissues. We found that overall, MyD88/methER-α interaction was detected in 68/190 of tissue sections (Table 1). However, when tissue samples were segregated according to sex, a male-to-female bias emerged (57/103 vs 11/87). This bias is consistent with our data showing that estrogen disrupts the MyD88/methER-α complex. We then reasoned that since differences in estrogen production are most pronounced in gametogenic reproductive tissues, this bias should be more obvious in these tissues. Indeed, we found that whereas none (0/31) of ovarian tissues from women at non-menopausal age (range: 18-49) displayed MyD88/methER-α interactions, all testicular tissues tested (35/35) exhibited strong PLA staining (Table 1). The cells that stained positive in the testes were histologically identified as Leydig cells (Figure 3). This could be explained by the fact that Leydig cells do not have functional estrogen since they express estrogen sulfotransferase, an enzyme that modifies this hormone, rendering it unable to bind to its receptors (26). We further reasoned that if non-menopaused ovarian tissues do not have MyD88/methER-α complexes because they express high levels of estrogen, ovarian tissues from women at menopausal age, which are known to produce reduced levels of estrogen (27), should have MyD88/methER-α complexes. Indeed, we found that 5/17 ovarian samples from women aged 50-69 were positive for MyD88/methER-α PLA staining (Table 1 and Figure 3). Collectively, these data show a strong segregation of MyD88/methER-α interaction according to gender in gametogenic reproductive organs and suggest a role for estrogen in disrupting this complex *in situ*.

**Figure 3:**
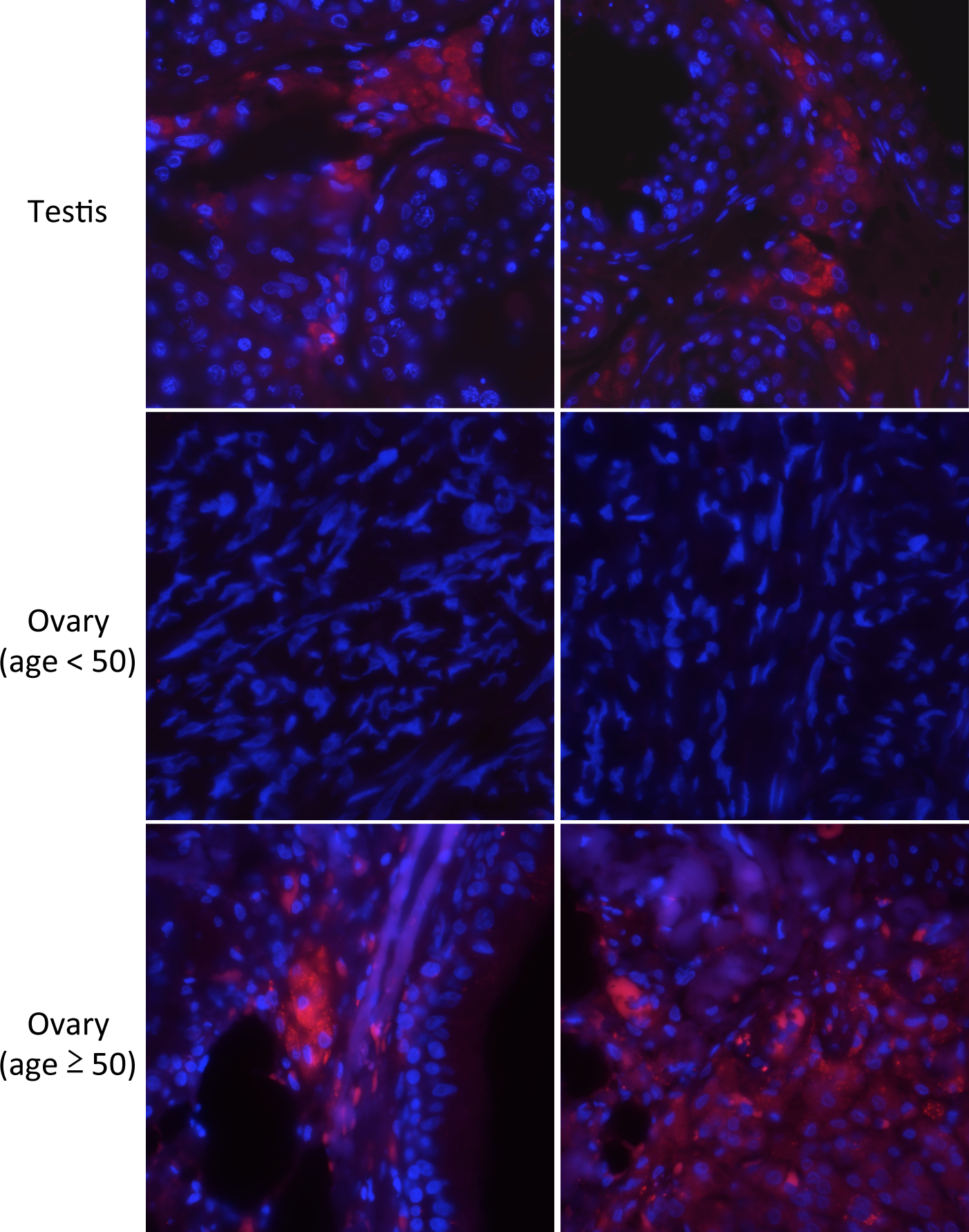
Segregation of MyD88/ER-α interaction according to gender. MyD88 and methER-α interaction in human normal testis and ovary tissues revealed by proximity ligation assay (red dots). Magnification: 60x

**Table 1:** MyD88-methER-α interaction in normal human tissue.

|  | Gender | Positive | Negative | Total | Percent |
| --- | --- | --- | --- | --- | --- |
| Overall total | Male + Female | 68 | 122 | 190 | 36 |
| Total male | Male | 57 | 46 | 103 | 55 |
| Total female | Female | 11 | 76 | 87 | 13 |
| Non-reproductive tissue* | Male | 22 | 46 | 68 | 32 |
|  | Female | 6 | 33 | 39 | 11 |
| Reproductive tissue** | Male | 35 | 0 | 35 | 100 |
|  | Female (age 18-49) | 0 | 31 | 31 | 0 |
|  | Female (age 50-70)*** | 5 | 12 | 17 | 29 |

\*\* Male: testis; Female: ovary
\*\*\* Except one 43 year-old female clinically determined to be menopausal

* Stomach, colon, bone, rectum, small intestine, esophagus, breast, liver, lung, spleen, kidney, lymph nodes, skin, thymus

** Male: testis; Female: ovary

*** Except one 43 yearCold female clinically determined to be menopaused

Sex-based differences in susceptibility to a wide variety of diseases are increasingly recognized as a confounding factor in the interpretation of clinical trials and responses to medication, with consequences on therapeutic decisions (28). Of the multiple biological parameters implicated in gender-based differences in inflammation, the hormone estrogen has emerged as a major contributing factor to the relative protection from inflammatory diseases observed in females in numerous experimental and epidemiological studies (1-12). Estrogen’s action in dampening inflammation seems to mainly implicate one of its receptors, ER-α, which exerts an inhibitory effect on NF-kB activation (14, 15, 29-31). These are hallmarks of the genomic pathway of ER-α function. Our data, on the other hand, implicate the methylated, cytoplasmic form of ER-α in gender disparity in response to inflammatory mediators. This pathway allows a rapid response to infection or tissue injury and thus may be useful in the early stages of the insult. Interestingly, in our hands this response does not seem to be restricted to professional cells of innate immunity such as macrophages, but it also takes place in epithelial and interstitial cells. This is consistent with recent data indicating that these cell types in the colon, lung, liver, and testis express TLRs and/or IL-1R and respond to ligands by producing cytokines and chemokines (32-35).

In summary, our results indicate that gender bias in inflammatory responses may be at least partly due to the MyD88/methER-α complex formation and its regulation by estrogen.

## Author contributions

RES, MB, KLC, KL, AK, NH, and FV conducted the experiments. BB, MLR, SL, and SM, provided critical insights and/or designed the experiments. IT selected and readpathology slides. TR wrote the manuscript. IC and TR supervised the work.

## Acknowledgements

This work was supported by the Ligue Contre le Cancer, the Institut National du Cancer (INCa), and the Integrated Research Site of Lyon (LYric).

